# RGT: a toolbox for the integrative analysis of high throughput regulatory genomics data

**DOI:** 10.1101/2022.12.31.522372

**Authors:** Zhijian Li, Chao-Chung Kuo, Fabio Ticconi, Mina Shaigan, Eduardo Gade Gusmao, Manuel Allhoff, Martin Manolov, Martin Zenke, Ivan G. Costa

**Affiliations:** Institute for Computational Genomics, RWTH Aachen University, Medical Faculty, 52074 Aachen, Germany; Joint Research Center for Computational Biomedicine, RWTH Aachen University Hospital, 52074 Aachen, Germany; Department of Cell Biology, Institute of Biomedical Engineering, RWTH Aachen University Medical School, 52074 Aachen, Germany; Helmholtz Institute for Biomedical Engineering, RWTH Aachen University, 52074 Aachen, Germany; Department of Hematology, Oncology, Hemostaseology, and Stem Cell Transplantation, Faculty of Medicine, RWTH Aachen University, 52074 Aachen, Germany

**Keywords:** Regulatory genomics, Motif analysis, Intersection algebra, Visualization, Footprinting, Differential peaks

## Abstract

**Background:** Massive amounts of data are produced by combining next-generation sequencing (NGS) with complex biochemistry techniques to characterize regulatory genomics profiles, such as protein-DNA interaction and chromatin accessibility. Interpretation of such high-throughput data typically requires different computation methods. However, existing tools are usually developed for a specific task, which makes it challenging to analyze the data in an integrative manner.

**Results:** We here describe the Regulatory Genomics Toolbox (RGT), a computational library for the integrative analysis of regulatory genomics data. RGT provides different functionalities to handle genomic signals and regions. Based on that, we developed several tools to perform distinct downstream analyses, including the prediction of transcription factor binding sites using ATAC-seq data, identification of differential peaks from ChIP-seq data, and detection of triple helix mediated RNA and DNA interactions, visualization, and finding an association between distinct regulatory factors.

**Conclusion:** We present here RGT; a framework to facilitate the customization of computational methods to analyze genomic data for specific regulatory genomics problems. RGT is a comprehensive and flexible Python package for analyzing high throughput regulatory genomics data and is available at: https://github.com/CostaLab/reg-gen. The documentation is available at: https://reg-gen.readthedocs.io

## Background

The combination of next-generation sequencing (NGS) with complex biochemistry techniques enables profiling of distinct epigenetic and regulatory features of cells in a genome-wide manner. Two examples are chromatin immunoprecipitation followed by sequencing (ChIP-seq) for protein-DNA interaction [1] and assay for transposase-accessible chromatin using sequencing (ATAC-seq) for open chromatin [2]. These techniques allow the studying of epigenetic dynamics in cellular processes such as cell differentiation [3, 4] and the characterization of the regulatory landscape of diseases such as human cancers [5]. Analysis of such data typically requires multistep computational pipelines that usually include:

- low-level methods (read alignment, quality control),
- medium-level methods for detection of genomic regions with relevant epigenetic signals (processing of genomic profiles, peak calling, differential peak calling, computational footprinting), and
- high-level methods for visual representation and integrative analysis with further genomic data (association with gene expression and further epigenetic data, detection of transcription factor binding sites, and functional enrichment analysis).

Figure 1 gives an example of a common ChIP-seq data analysis pipeline. It includes on the low level the use of a read aligner, such as BWA [6]; on the medium level a peak calling method, such as MACS2 [7], for the detection of regions with the presence of potential protein-DNA interactions; and on the high level a motif match procedure, such as FIMO [8], to find transcription factor binding sites inside peaks as well as R functions for the visualization of genomic signals, such as Genomics Ranges [9]. A similar pipeline for ATAC-seq data analysis is described in Supplementary Figure 1.

**Figure 1.**
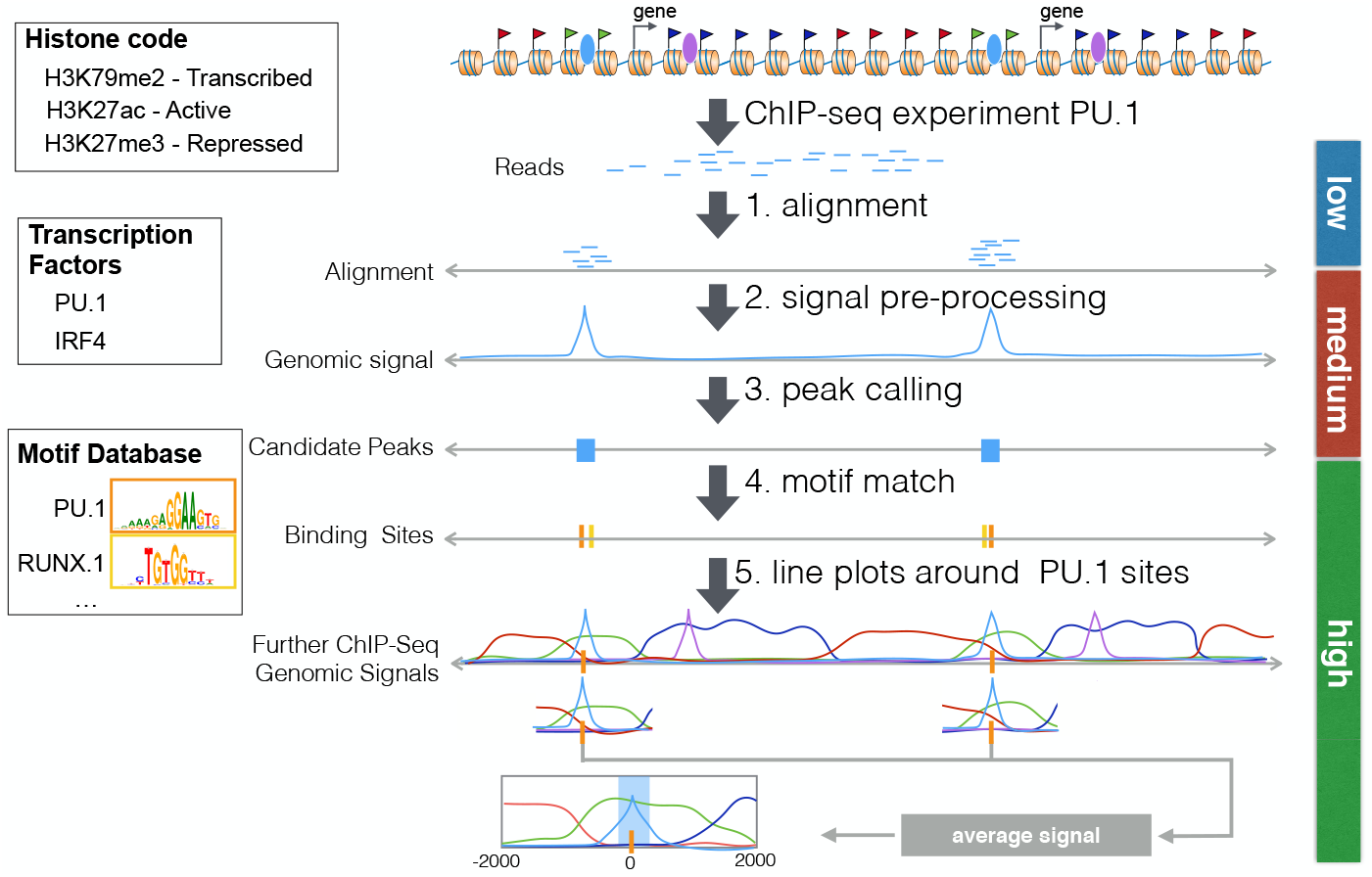
Example of a typical pipeline for the analysis of a transcription factor ChlP-seq experiment. First, the reads are aligned to the genome (step 1, low-level analysis). A peak caller receives these aligned reads as input and typically creates an intermediary representation called *genomic signal*. Based on this genomic signal, the peak caller then detects regions with a higher value than the background. These candidate peaks represent the regions with DNA-protein interaction sites (steps 2 and 3, medium level). Several downstream analyses are then performed, such as the detection of motif-predicted binding sites inside the peaks (step 4, high-level analysis) or line plots displaying average genomic signals of other ChIP-seq experiments around the predicted peaks or binding sites (step 5, high-level analysis).

The definition of analysis pipelines depends on the biological study as well as on the used NGS technique. Its complexity, which includes the use of several bioinformatics tools that may require command-line usage and/or scripting skills, makes the analysis of epigenomics data so far inaccessible for non-bioinformatics specialists. Moreover, the development of bioinformatics tools for medium-level analysis needs to take into account specific characteristics of the used NGS protocols [10, 11].

For example, ChIP-seq experiments require the computational estimation of the read extension sizes [11]. It also requires a signal correction with control experiments, as the local chromatin structure may influence the ChIP-seq signal [12]. In contrast, footprint analysis of ATAC-seq data does not require the estimation of read extension sizes, as the start of the read corresponds to the cleavage position. However, ATAC-seq analysis demands the correction of Tn5 cleavage bias [13]. Moreover, some aspects, such as PCR amplification artifacts, are shared by ChIP-seq and ATAC-seq experiments [11]. Clearly, the development of tools for the analysis of epigenetic data is greatly facilitated by a flexible and easy-to-handle computational library. This library should support genomic data I/O as well as usual pre-processing methods, such as fragment size estimation and the correction of sequence bias. Regarding high-level tasks, the library should provide structure to allow sequence analysis (i.e. motif matching), interval algebra (i.e. measuring overlap between peaks), or associating signals with regions (i.e. line plots showing signal strength around peaks).

## Implementation

We developed RGT in Python by following the object-oriented approach. The core classes provide functionalities for handling data structures that are related to questions about regulatory genomics. Based on the cores, we implemented several computational tools to perform various downstream analyses (Fig. 2). These include previously described HINT tools for ATAC-seq/DNase-seq footprinting [13–15], the differential peak caller THOR [16] and a library to characterize triple helix mediated RNA-DNA interactions [17]. RGT also includes some functionalities such as motif binding sites prediction and enrichment analysis (Motif Analysis), as well as methods for association and visualization of genomic signals (RGT-Viz). We describe below the basic structures and the novel Motif Analysis and the RGT-Viz frameworks.

**Figure 2.**
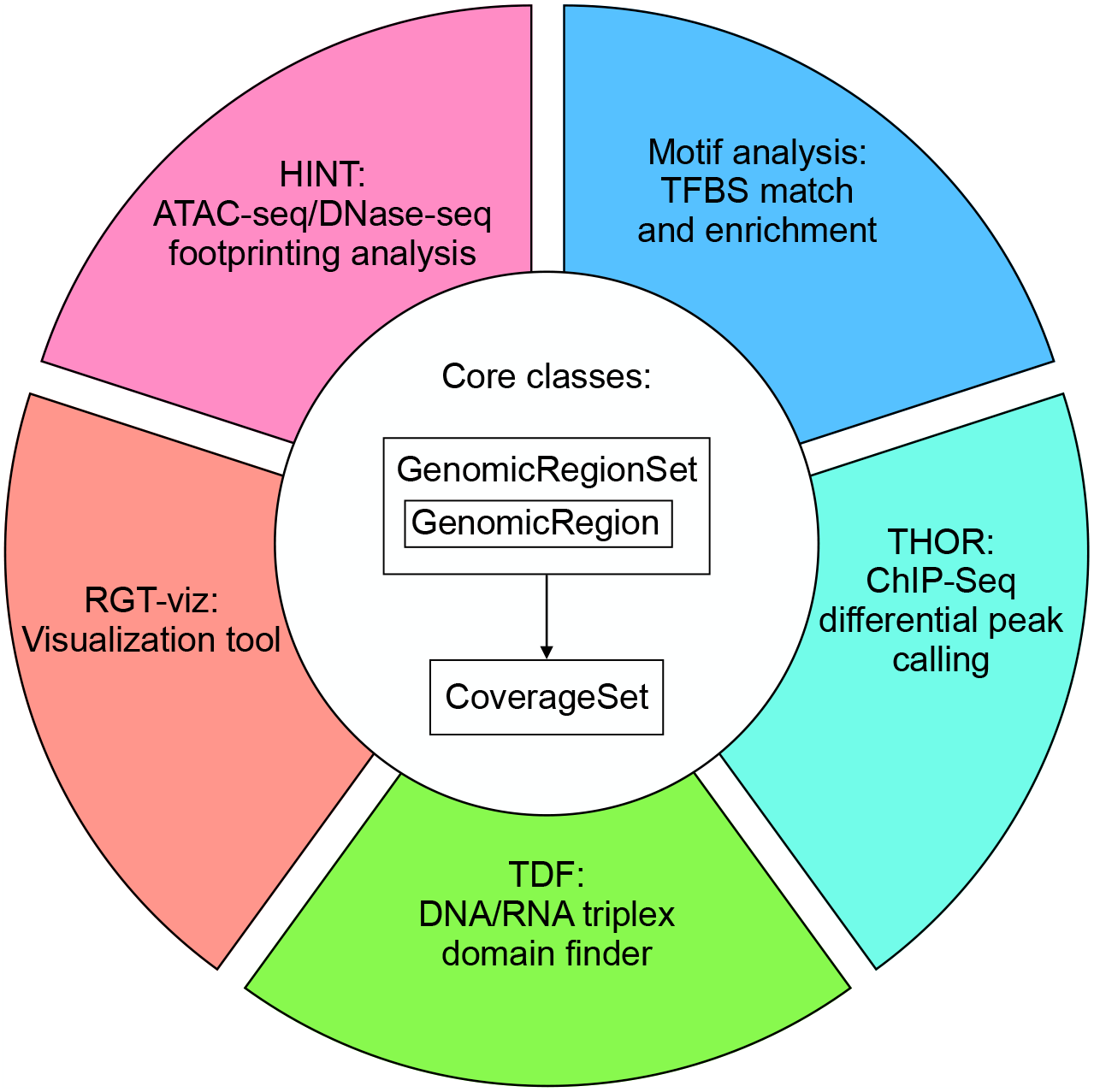
Overview of RGT core classes and tools. RGT provides three core classes to handle the genomic regions and signals. Each genomic region is represented by GenomicRegion class and multiple regions are represented by GenomicRegionSet class. The genomic signals are represented CoverageSet class. These classes serve as the core data structures of RGT for handling genomic regions and signals. Based on these classes, we developed several tools for analyzing regulatory genomics data as represented by different colors, namely, HINT for footprinting analysis of ATAC/DNase-seq data; RGT-viz for finding associations between chromatin experiments; TDF for DNA/RNA triplex domain finder; THOR for differential peak calling of ChIP-seq data; Motif analysis for transcription factor binding sites matching and enrichment.

### Core classes

Analysis of high-throughput regulatory genomics data is mostly based on the manipulation of two common data structures: genomic signals which represent the abundance of sequencing reads on the genome and genomic regions which represents candidate regions. In RGT, we implemented three classes, i.e., GenomicRegion, GenomicRegionSet, and CoverageSet, to represent a single region, multiple regions, and genomic signals, respectively. In each of the classes, we implemented several functions to perform basic data processing. For example, CoverageSet provides functions for fragment extension estimation, signal smoothing, GC-content bias correction, and input DNA normalization. These procedures are crucial for the particular downstream analysis of chromatin sequencing data, such as peak calling and footprinting. For computational efficiency, functions related to GenomicRe-gionsSet and interval-related algebra have been implemented in C. Moreover, RGT contains I/O functions of common genomic file formats such as Binary Alignment Map (BAM) files for alignments of reads, (big)wig files for genomic profiles, and bed files for genomic regions by exploring pysam [18, 19] related functions.

These core classes provide a powerful infrastructure for the development of methods dealing with regulatory genomics data. As an example of the simplicity, versatility, and power of RGT, we include a tutorial on how to build a simple peak caller with less than 50 lines of codes: https://reg-gen.readthedocs.io/en/latest/rgt/tutorial-peak-calling.html.

### Finding associations between chromatin experiments with RGT-viz

A typical problem in regulatory genomics is to associate results of distinct experiments, i.e. overlap between distinct histone marks or a given histone mark in distinct cells. RGT-viz provides a collection of statistical tests and tools for the association and visualization of genomic data such as genomic regions and genomic signals (Fig. 3a).

**Figure 3.**
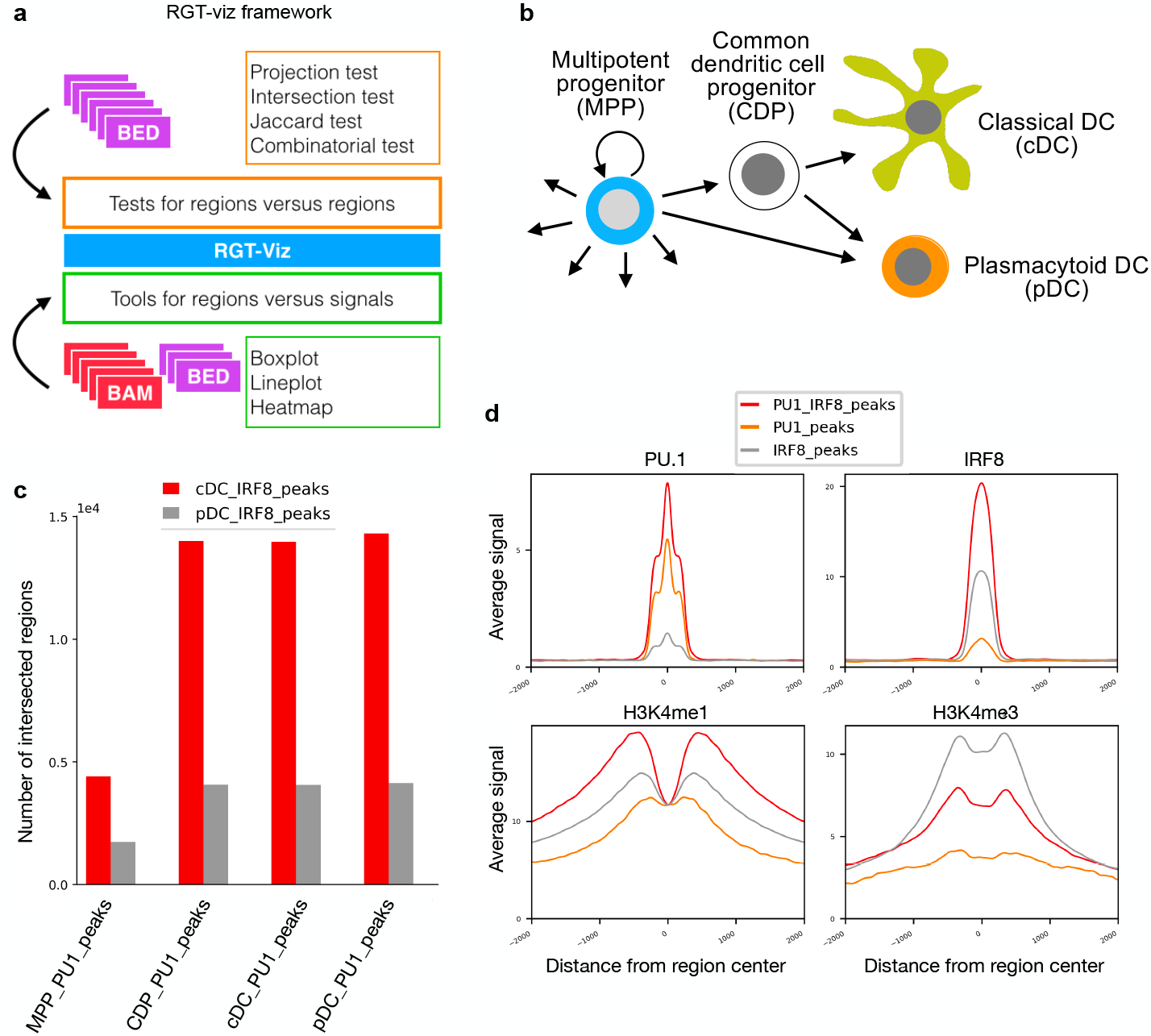
Overview and case study of RGT-viz. **a,** RGT-viz provides several tests for regions versus regions and visualization tools for regions versus signals by taking BED and BAM files as input. **b,** Dendritic cell development. DCs develop from multipotent progenitors (MPPs), which commit into DC-restricted common dendritic cell progenitors (CDPs). CDPs differentiate into classical DCs (cDCs) and plasmacytoid DCs (pDCs). **c,** Intersection test shows that the IRF8 binding sites in cDC and pDC are associated with the PU.1 binding sites in MPP, CDP, cDC, and pDC. **d,** Line plots showing genomic signals of different histone modifications on the PU.1/IRF8 peaks in cDC.

In the tests of regions versus regions, a set of reference and query regions, both in BED format, are required as inputs. The aim is to evaluate the association between the reference and the query. For this, RGT-viz provides the following tests:

- Projection test: This test compares a query set, i.e. ChIP-seq of transcription factors with a larger reference set, i.e. ChIP-seq peaks of a regulatory region (H3K4me3 or H3K4me1 marks). It estimates the overlap of the query to the reference and contrasts with the coverage of the reference in the complete genome. A binomial test is then used to indicate if the coverage of the query in the reference is higher than the reference of the reference to the genome [20] (Supplementary Fig. 2a).
- Intersection Test: This test is based on measuring the intersection between a pair of genomic regions and comparing it to the expected intersection on random region sets. Random regions are obtained by evaluating permutations (with size equal to the input regions) of the union of regions in the pair of queries [21] (Supplementary Fig. 2b). The statistical test is based on empirical *p*—values.
- Combinatorial Test: The combinatorial test is appropriate for two-way comparisons. For example, you want to check the proportion of peaks of two (or more) transcription factors on two (or more) cell types. For this, it creates a background distribution per reference sets (cells) by considering the union of all query sets (TFs) in that cell. It then creates count statistics per cell and compares if the number of binding sites in a cell for a given TF is higher than in another cell by using a Chi-squared test (Supplementary Fig. 2c).
- Jaccard Measure: This measures the amount of overlap between the reference and the query using the Jaccard index (also called Jaccard similarity coefficient) [22]. Given two region sets A and B, it measures the ratio of intersecting base pairs in relation to the regions associated with the union of A and B. Through this Jaccard index, the amount of intersection can be expressed by a value between zero to one (Supplementary Fig. 2d). This test explores a randomization approach, i.e. random selection of genomic region sets with the same number/size regions, to estimate empirical p—values.

Another important functionality is the visualization of distinct genomic signals, as described below. To visualize the signals in different regions, the following tools are provided:

- Boxplot: It compares the number of fragments from different ChIP-seq experiments on the given region set. This can be used for example to contrast the signal of distinct ChIP-seq TFs over promoter regions (H3K4me3 peaks). Conceptually, the generation of a boxplot is simply counting the number of reads within the region set and then plotting these counts in boxplot (Supplementary Fig. 3a). RGT-Viz provides functionalities to normalize the individual libraries regarding library sizes.
- Lineplot and heatmap: Line plot and heatmap are used to display the distribution of reads within a given region set. Specifically, each region is first extended with the given window size which defines the boundaries for plotting. Next, the coverage of reads on the given regions is calculated based on the given bin size and step size. Finally, the line plot or heatmap is generated. The line plot shows average signals over all regions in the region set while the heatmap displays the signals of all regions (Supplementary Fig. 3b-c).

We here provided a case study using RGT-viz to investigate dendritic cell (DC) development (Fig. 3b). We collected ChIP-seq data of the transcription factors PU.1 and IRF8, and five histone modifications (i.e., H3K4me1, H3K4me3, H3K9me3, H3K27me3, and H3K27ac) for each of the cell types [4, 16, 23, 24] (Supplementary Table 1). PU.1 is one of the master regulators of hematopoiesis and is expressed by all hematopoietic cells [25] and IRF8 is believed to co-bind with PU.1 to control the differentiation of DC progenitors (DCP) towards specific DC sub-types [4, 26]. We mapped the sequencing reads to mm9 using BWA [6] and called the peaks with MACS2 [7].

We performed an intersection test between PU.1 and IRF8 peaks from different cell types to check for if PU.1 and IRF8 co-binding during DC differentiation. Of note, IRF8 ChIP-seq only detected peaks in classic and plasmacytoid DC (cDC and pDC, respectively), as this TF is not expressed in multipotent DC (MPP) and expressed only at low levels in common DC progenitors (CDP).

This test reveals that PU.1 and IRF8 are significantly associated in all cell types, while the co-binding was two times higher as measured by χ^2^ statistics in cDCs than pDCs. Moreover, we observed that a high overlap of binding sites of cDC IRF8 peaks is already quite high with CDP PU.1 peaks. This indicates that PU.1 binding prepares the chromatin for IRF8 binding already CDP, showing DC priming in CDP (Fig. 3c and Supp. Table 2).

We next asked if the co-binding regions are associated with different regulatory regions (enhancers vs. promoters). For this, we defined the set of peaks with both PU.1. and IRF8 binding, or only with PU.1. or only IRF8 binding in cDC and pDC cells by using intersect and subtracting functions from the core class GenomicRegionSet of RGT. We then generated line plots of PU.1, IRF8, H3K4me1, and H3K4me3 on these three sets of regions in cDC (Figure 3d). We observed that peaks with PU.1-IRF8 co-binding have higher ChIP-seq peaks for either factor indicating that co-binding strengthens the binding affinity of both TFs. Moreover, H3K4me1 signals are strong for PU.1 and IRF8 co-binding, while IRF8 only has stronger H3K4me3 marks. This suggests an association of PU.1 and IRF8 co-binding with enhancers, while IRF8 exclusive binding is more associated with promoters. These examples demonstrate how RGT-Viz can be used to explore associations and interpretation of genomic data.

### Transcription factor motif matching and enrichment with motif analysis

Motif analysis is a framework to perform transcription factor motif matching and motif enrichment. Motif matching aims to find transcription factor binding sites (TFBSs) for a set of TFs in a set of genomic regions of interest (Figure 4a). For this, RGT has its own class, i.e., MotifSet for storing TF motifs from known repositories, such as UniPROBE [27], JASPAR [28] and HOCOMOCO [29]. In addition, users are also allowed to add new motif repositories. RGT uses an efficient Motif Occurrence Detection Suite (MOODS) algorithm to find binding site locations and bit-scores [30]. Note that MOODS was originally implemented in C++ and we have adapted it to a Python package (https://pypi.org/project/MOODS-python/). Next, RGT uses a dynamic programming algorithm [31] to determine a bit-score cut-off threshold based on the false positive rate of 10^-4^. The predicted binding sites can be obtained with p-values between 10^-5^ to 10^-3^.

**Figure 4.**
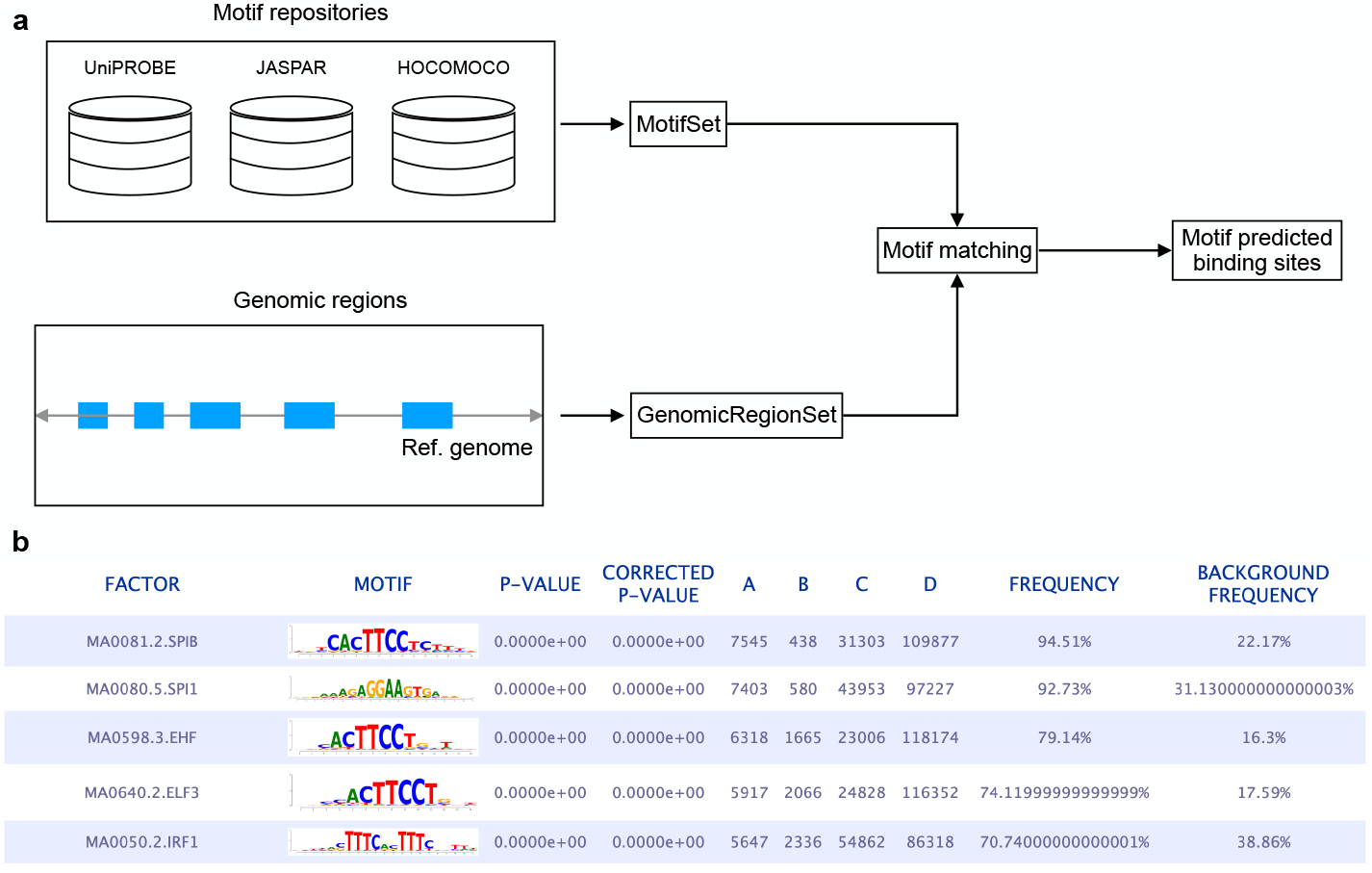
Schematics of motif matching and enrichment analysis. **a,** Motif matching detects binding sites for a set of TFs against multiple genomic regions. The motifs were collected from public repositories such as UniPROBE, JASPAR, and HOCOMOCO. The position weight matrix (PWM) for each TF is used to calculate a binding affinity score per position. The genomic regions are usually obtained by peak calling based on ChIP-seq or ATAC-seq data. **b,** Screenshot showing the top 5 TFs identified by motif enrichment analysis from the overlapping peaks between PU.1 and IRF8 in cDC cells.

The motif enrichment module evaluates which transcription factors are more likely to occur in certain genomic regions than in “background regions” based on the motif-predicted binding sites (MPBS) from motif matching (Figure 4b). To determine the significance, we performed Fisher’s exact test for each transcription factor and corrected the p-values with the Benjamini-Hochberg procedure. More specifically, we provided three types of tests:

- Input regions vs. Background regions: In this test, all input regions are verified against background regions that are either user-provided or randomly generated with the same average length distribution as the original input regions.
- Gene-associated regions vs. Non-gene-associated regions: In this test, we would like to check whether a group of regions that are associated with genes of interest (e.g. up-regulated genes) is enriched for some transcription factors vs. regions that are not associated with those genes. The input regions are divided into two groups by performing gene-region association that considers promoter-proximal regions, gene body, and distal regions. After the association, we perform a Fisher’s exact test followed by multiple testing corrections as mentioned in the previous analysis type.
- Promoter regions of input genes vs. Background regions: In this test, we take all provided genes, find their promoter regions in the target organism, and create a “target regions” BED file from those. A background file is created by using the promoter regions of all genes not included in the provided gene list. Next, motif matching is performed on the target and background regions and a Fisher’s exact test is executed.

Finally, the enrichment regions are provided in an HTML interface. An example of motif enrichment analysis on the PU.1 and IRF8 co-binding peaks in cDC is provided in Fig. 4b). We observed that PU.1 (and ETS family) motifs were ranked at the top and an IRF family motif at fifth (IRF1; MA0050.2.IRF1). This again demonstrates how Motif Matching can recover expected regulatory players from regulatory sequences.

### Additional Tools based on RGT

Several additional tools that explored and extended classes from RGT to tackle specific regulatory genomics problems are available. HINT is a framework that uses open chromatin data to identify the active transcription factor binding sites (TFBS). We originally developed this method for DNase-seq data [14, 15] and later extended it to ATAC-seq data by taking the protocol-specific artifacts into account [13]. Footprint analysis requires base pair resolution signals in contrast to peak calling problems, which are based on signals on windows with more than 50bps. Therefore, HINT has a GenomicSignal class, which deals with ATAC-seq, and DNA-seq signals such as cleavage bias correction, base pair counting, and signal smoothing. Moreover, HINT makes use of the previously described motif-matching functionality provided by RGT to characterize motifs related to ATAC-seq footprints. These can be explored in differential footprinting analysis to detect relevant TFs associated with different biological conditions. This method has been widely used to study, among others, cell differentiation [13, 32] and diseases [33–36].

THOR is a Hidden Markov Model-based approach to detect and analyze differential peaks in two sets of ChIP-seq data from distinct biological conditions with replicates [16]. As a first step, THOR needs to create and normalize ChIP-seq signals from distinct experiments. Among others, THOR extended functionalities of the base class CoverageSet to a MultipleCoverageSet class to deal with multiple signals at a time and to provide global normalization methods, such as trimmed means of M-values (TMM). Finally, Triplex Domain Finder (TDF) characterizes the triplex-forming potential between RNA and DNA regions [17]. TDF explores functionality provided by RGT/RGT-viz to build statistical tests for characterizing DNA binding domains in lncRNAs.

## Discussion

We here presented the regulatory genomics toolbox (RGT), a versatile toolbox for the analysis of high-throughput regulatory genomics data. RGT was programmed in an oriented-object fashion and its core classes provided functionalities to handle typical regulatory genomics data: regions and signals. Based on these core classes, RGT built distinct regulatory genomics tools, i.e., HINT for footprinting analysis, TDF for finding DNA-RNA triplex, THOR for ChIP-seq differential peak calling, motif analysis for TFBS matching and enrichment, and RGT-viz for regions association tests and data visualization. These tools have been used in several of epigenomics and regulatory genomics works to study cell differentiation and regulation [32, 35, 37–42]. We envision that RGT can facilitate the development of computational methods for the analysis of high-throughput regulatory genomics data as a powerful and flexible framework in the future.

## Acknowledgements

This project was funded by the German Research Foundation (DFG) and the E: MED Consortia Fibromap funded by the German Ministry of Education and Science (BMBF).

## Author’s contributions

ZL, CCK, FT, EGG, MA, and MS developed RGT under the supervision of IC. MM supported the RGT documentation. MZ interpreted the data related to DC development. ZL and CCK jointly wrote the paper with revisions from IGC. MZ contributed to the RGT-Viz concept and case studies. All authors read and approved the final manuscript.

## Code availability

RGT is available as an open-source Python package at GitHub: https://github.com/CostaLab/reg-gen. The documentation including methods description and tools tutorial is available in: https://reg-gen.readthedocs.io.

## Additional Files

Additional file 1: Supplementary figures

## References

1. Park, P.J.: ChIP-seq: advantages and challenges of a maturing technology. Nature Reviews Genetics 10(10), 669–680 (2009)

2. Buenrostro, J.D., Giresi, P.G., Zaba, L.C., Chang, H.Y., Greenleaf, W.J.: Transposition of native chromatin for fast and sensitive epigenomic profiling of open chromatin, DNA-binding proteins and nucleosome position. Nature methods 10(12), 1213–1218 (2013)

3. Lara-Astiaso, D., Weiner, A., Lorenzo-Vivas, E., Zaretsky, I., Jaitin, D.A., David, E., Keren-Shaul, H., Mildner, A., Winter, D., Jung, S., et al.: Chromatin state dynamics during blood formation. science 345(6199), 943–949 (2014)

4. Lin, Q., Chauvistré, H., Costa, I.G., Gusmao, E.G., Mitzka, S., Hänzelmann, S., Baying, B., Klisch, T., Moriggl, R., Hennuy, B., et al.: Epigenetic program and transcription factor circuitry of dendritic cell development. Nucleic acids research 43(20), 9680–9693 (2015)

5. Corces, M.R., Granja, J.M., Shams, S., Louie, B.H., Seoane, J.A., Zhou, W., Silva, T.C., Groeneveld, C., Wong, C.K., Cho, S.W., et al.: The chromatin accessibility landscape of primary human cancers. Science 362(6413), 1898 (2018)

6. Li, H., Durbin, R.: Fast and accurate long-read alignment with burrows–wheeler transform. Bioinformatics 26(5), 589–595 (2010)

7. Zhang, Z.D., Rozowsky, J., Snyder, M., Chang, J., Gerstein, M.: Modeling chip sequencing in silico with applications. PLoS Comput Biol 4(8), 1000158 (2008)

8. Grant, C.E., Bailey, T.L., Noble, W.S.: FIMO: scanning for occurrences of a given motif. Bioinformatics 27(7), 1017–1018 (2011)

9. Lawrence, M., Huber, W., Pages, H., Aboyoun, P., Carlson, M., Gentleman, R., Morgan, M.T., Carey, V.J.: Software for computing and annotating genomic ranges. PLoS computational biology 9(8), 1003118 (2013)

10. Furey, T.S.: ChIP-seq and beyond: new and improved methodologies to detect and characterize protein-DNA interactions. Nat Rev Genet 13(12), 840–852 (2012)

11. Meyer, C.A., Liu, X.S.: Identifying and mitigating bias in next-generation sequencing methods for chromatin biology. Nature Reviews Genetics 15(11), 709–721 (2014)

12. Diaz, A., Park, K., Lim, D.A., Song, J.S.: Normalization, bias correction, and peak calling for ChIP-seq. Statistical applications in genetics and molecular biology 11(3) (2012)

13. Li, Z., Schulz, M.H., Look, T., Begemann, M., Zenke, M., Costa, I.G.: Identification of transcription factor binding sites using ATAC-seq. Genome biology 20(1), 1–21 (2019)

14. Gusmao, E.G., Dieterich, C., Zenke, M., Costa, I.G.: Detection of active transcription factor binding sites with the combination of DNase hypersensitivity and histone modifications. Bioinformatics 30(22), 3143–3151 (2014)

15. Gusmao, E.G., Allhoff, M., Zenke, M., Costa, I.G.: Analysis of computational footprinting methods for DNase sequencing experiments. Nature methods 13(4), 303–309 (2016)

16. Allhoff, M., Seré, K., F. Pires, J., Zenke, M., G. Costa, I.: Differential peak calling of ChIP-seq signals with replicates with THOR. Nucleic acids research 44(20), 153–153 (2016)

17. Kuo, C.-C., Hänzelmann, S., Sentürk Cetin, N., Frank, S., Zajzon, B., Derks, J.-P., Akhade, V.S., Ahuja, G., Kanduri, C., Grummt, I., et al.: Detection of RNA–DNA binding sites in long noncoding RNAs. Nucleic acids research 47(6), 32–32 (2019)

18. Bonfield, J.K., Marshall, J., Danecek, P., Li, H., Ohan, V., Whitwham, A., Keane, T., Davies, R.M.: Htslib: C library for reading/writing high-throughput sequencing data. Gigascience 10(2), 007 (2021)

19. Danecek, P., Bonfield, J.K., Liddle, J., Marshall, J., Ohan, V., Pollard, M.O., Whitwham, A., Keane, T., McCarthy, S.A., Davies, R.M., et al.: Twelve years of samtools and bcftools. Gigascience 10(2), 008 (2021)

20. Favorov, A., Mularoni, L., Cope, L.M., Medvedeva, Y., Mironov, A.A., Makeev, V.J., Wheelan, S.J.: Exploring massive, genome scale datasets with the genometricorr package. PLoS computational biology 8(5), 1002529 (2012)

21. Pape, U.J., Klein, H., Vingron, M.: Statistical detection of cooperative transcription factors with similarity adjustment. Bioinformatics 25(16), 2103–2109 (2009)

22. Real, R., Vargas, J.M.: The probabilistic basis of jaccard’s index of similarity. Systematic biology 45(3), 380–385 (1996)

23. Allhoff, M., Seré, K., Chauvistré, H., Lin, Q., Zenke, M., Costa, I.G.: Detecting differential peaks in chip-seq signals with odin. Bioinformatics 30(24), 3467–3475 (2014)

24. Xu, H., Li, Z., Kuo, C.-C., Götz, K., Look, T., de Toledo, M.A.S., Seré, K., Costa, I.G., Zenke, M.: A lncrna identifies irf8 enhancer element in negative feedback control of dendritic cell differentiation. bioRxiv (2022)

25. Belz, G.T., Nutt, S.L.: Transcriptional programming of the dendritic cell network. Nature Reviews Immunology 12(2), 101–113 (2012)

26. Chauvistré, H., Küstermann, C., Rehage, N., Klisch, T., Mitzka, S., Felker, P., Rose-John, S., Zenke, M., Seré, K.M.: Dendritic cell development requires histone deacetylase activity. European journal of immunology 44(8), 2478–2488 (2014)

27. Newburger, D.E., Bulyk, M.L.: Uniprobe: an online database of protein binding microarray data on protein–dna interactions. Nucleic acids research 37(suppl_1), 77–82 (2009)

28. Castro-Mondragon, J.A., Riudavets-Puig, R., Rauluseviciute, I., Berhanu Lemma, R., Turchi, L., Blanc-Mathieu, R., Lucas, J., Boddie, P., Khan, A., Manosalva Pérez, N., et al.: JASPAR 2022: the 9th release of the open-access database of transcription factor binding profiles. Nucleic acids research 50(D1), 165–173 (2022)

29. Kulakovskiy, I.V., Vorontsov, I.E., Yevshin, I.S., Sharipov, R.N., Fedorova, A.D., Rumynskiy, E.I., Medvedeva, Y.A., Magana-Mora, A., Bajic, V.B., Papatsenko, D.A., et al.: HOCOMOCO: towards a complete collection of transcription factor binding models for human and mouse via large-scale ChIP-Seq analysis. Nucleic acids research 46(D1), 252–259 (2018)

30. Korhonen, J., Martinmäki, P., Pizzi, C., Rastas, P., Ukkonen, E.: MOODS: fast search for position weight matrix matches in DNA sequences. Bioinformatics 25(23), 3181–3182 (2009)

31. Wilczynski, B., Dojer, N., Patelak, M., Tiuryn, J.: Finding evolutionarily conserved cis-regulatory modules with a universal set of motifs. BMC bioinformatics 10(1), 1–11 (2009)

32. Heller, S., Li, Z., Lin, Q., Geusz, R., Breunig, M., Hohwieler, M., Zhang, X., Nair, G.G., Seufferlein, T., Hebrok, M., Sander, M., Julier, C., Kleger, A., Costa, I.G.: Transcriptional changes and the role of ONECUT1 in hPSC pancreatic differentiation. Communications Biology 4(1), 1–12 (2021). doi:10.1038/s42003-021-02818-3

33. Caldwell, A.B., Liu, Q., Schroth, G.P., Galasko, D.R., Yuan, S.H., Wagner, S.L., Subramaniam, S.: Dedifferentiation and neuronal repression define familial Alzheimer’s disease. Science advances 6(46), 5933 (2020)

34. Li, Z., Kuppe, C., Ziegler, S., Cheng, M., Kabgani, N., Menzel, S., Zenke, M., Kramann, R., Costa, I.G.: Chromatin-accessibility estimation from single-cell atac-seq data with scopen. Nature communications 12(1), 1–14 (2021)

35. Philippi, A., Heller, S., Costa, I.G., Senée, V., Breunig, M., Li, Z., Kwon, G., Russell, R., Illing, A., Lin, Q., Hohwieler, M., Degavre, A., Zalloua, P., Liebau, S., Schuster, M., Krumm, J., Zhang, X., Geusz, R., Benthuysen, J.R., Wang, A., Chiou, J., Gaulton, K., Neubauer, H., Simon, E., Klein, T., Wagner, M., Nair, G., Besse, C., Dandine-Roulland, C., Olaso, R., Deleuze, J.-F., Kuster, B., Hebrok, M., Seufferlein, T., Sander, M., Boehm, B.O., Oswald, F., Nicolino, M., Julier, C., Kleger, A.: Mutations and variants of ONECUT1 in diabetes. Nature Medicine 27(11), 1928–1940 (2021). doi:10.1038/s41591-021-01502-7

36. Kuppe, C., Ramirez Flores, R.O., Li, Z., Hayat, S., Levinson, R.T., Liao, X., Hannani, M.T., Tanevski, J., Wünnemann, F., Nagai, J.S., et al.: Spatial multi-omic map of human myocardial infarction. Nature, 1–12 (2022)

37. Barsoum, M., Stenzel, A.T., Bochyńska, A., Kuo, C.-C., Tsompanidis, A., Sayadi-Boroujeni, R., Bussmann, P., Lüscher-Firzlaff, J., Costa, I.G., Lüscher, B.: Loss of the Ash2l subunit of histone H3K4 methyltransferase complexes reduces chromatin accessibility at promoters. Scientific Reports 12(1), 21506 (2022). doi:10.1038/s41598-022-25881-0

38. Bentsen, M., Goymann, P., Schultheis, H., Klee, K., Petrova, A., Wiegandt, R., Fust, A., Preussner, J., Kuenne, C., Braun, T., Kim, J., Looso, M.: ATAC-seq footprinting unravels kinetics of transcription factor binding during zygotic genome activation. Nature Communications 11(1) (2020). doi:10.1038/s41467-020-18035-1

39. Greco, C.M., Koronowski, K.B., Smith, J.G., Shi, J., Kunderfranco, P., Carriero, R., Chen, S., Samad, M., Welz, P.-S., Zinna, V.M., Mortimer, T., Chun, S.K., Shimaji, K., Sato, T., Petrus, P., Kumar, A., Vaca-Dempere, M., Deryagin, O., Van, C., Kuhn, J.M.M., Lutter, D., Seldin, M.M., Masri, S., Li, W., Baldi, P., Dyar, K.A., Muñoz-Cáoves, P., Benitah, S.A., Sassone-Corsi, P.: Integration of feeding behavior by the liver circadian clock reveals network dependency of metabolic rhythms. Science Advances 7(39) (2021). doi:10.1126/sciadv.abi7828

40. Roquilly, A., Jacqueline, C., Davieau, M., Mollé, A., Sadek, A., Fourgeux, C., Rooze, P., Broquet, A., Misme-Aucouturier, B., Chaumette, T., Vourc’h, M., Cinotti, R., Marec, N., Gauttier, V., McWilliam, H.E.G., Altare, F., Poschmann, J., Villadangos, J.A., Asehnoune, K.: Alveolar macrophages are epigenetically altered after inflammation, leading to long-term lung immunoparalysis. Nature Immunology 21(6), 636–648 (2020). doi:10.1038/s41590-020-0673-x

41. Sentürk Cetin, N., Kuo, C.-C., Ribarska, T., Li, R., Costa, I.G., Grummt, I.: Isolation and genome-wide characterization of cellular DNA:RNA triplex structures. Nucleic Acids Research 47(5), 2306–2321 (2019). doi:10.1093/nar/gky1305

42. Willcockson, M.A., Healton, S.E., Weiss, C.N., Bartholdy, B.A., Botbol, Y., Mishra, L.N., Sidhwani, D.S., Wilson, T.J., Pinto, H.B., Maron, M.I., Skalina, K.A., Toro, L.N., Zhao, J., Lee, C.H., Hou, H., Yusufova, N., Meydan, C., Osunsade, A., David, Y., Cesarman, E., Melnick, A.M., Sidoli, S., Garcia, B.A., Edelmann, W., Macian, F., Skoultchi, A.I.: H1 histones control the epigenetic landscape by local chromatin compaction. Nature 589(7841), 293–298 (2021). doi:10.1038/s41586-020-3032-z

