## Supplemental Figures for "RGT: a toolbox for the integrative analysis of high throughput regulatory genomics data"

December 30, 2022

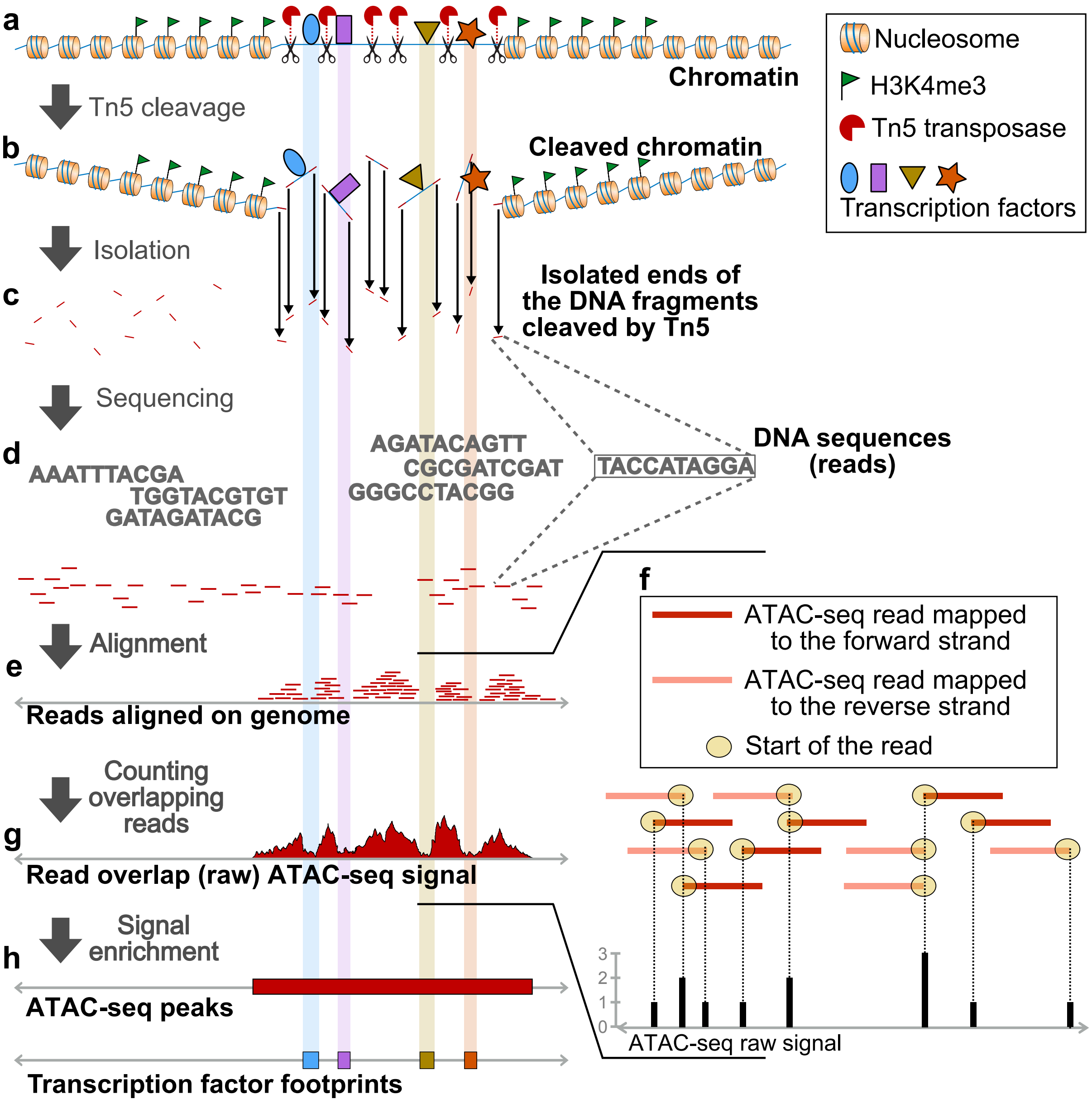


**Supplementary Fig. 1. An example pipeline for the analysis of ATAC-seq data.** Similar to ChIP-seq data, the first step after obtaining the sequencing data is to perform the alignment. Based on the aligned reads, a genome-wide signal is created by counting overlapping reads on the start site. Next, chromatin-accessible regions are detected by peak calling. Finally, transcription factor footprints are identified by searching for footprint-like Tn5 cleavage patterns.

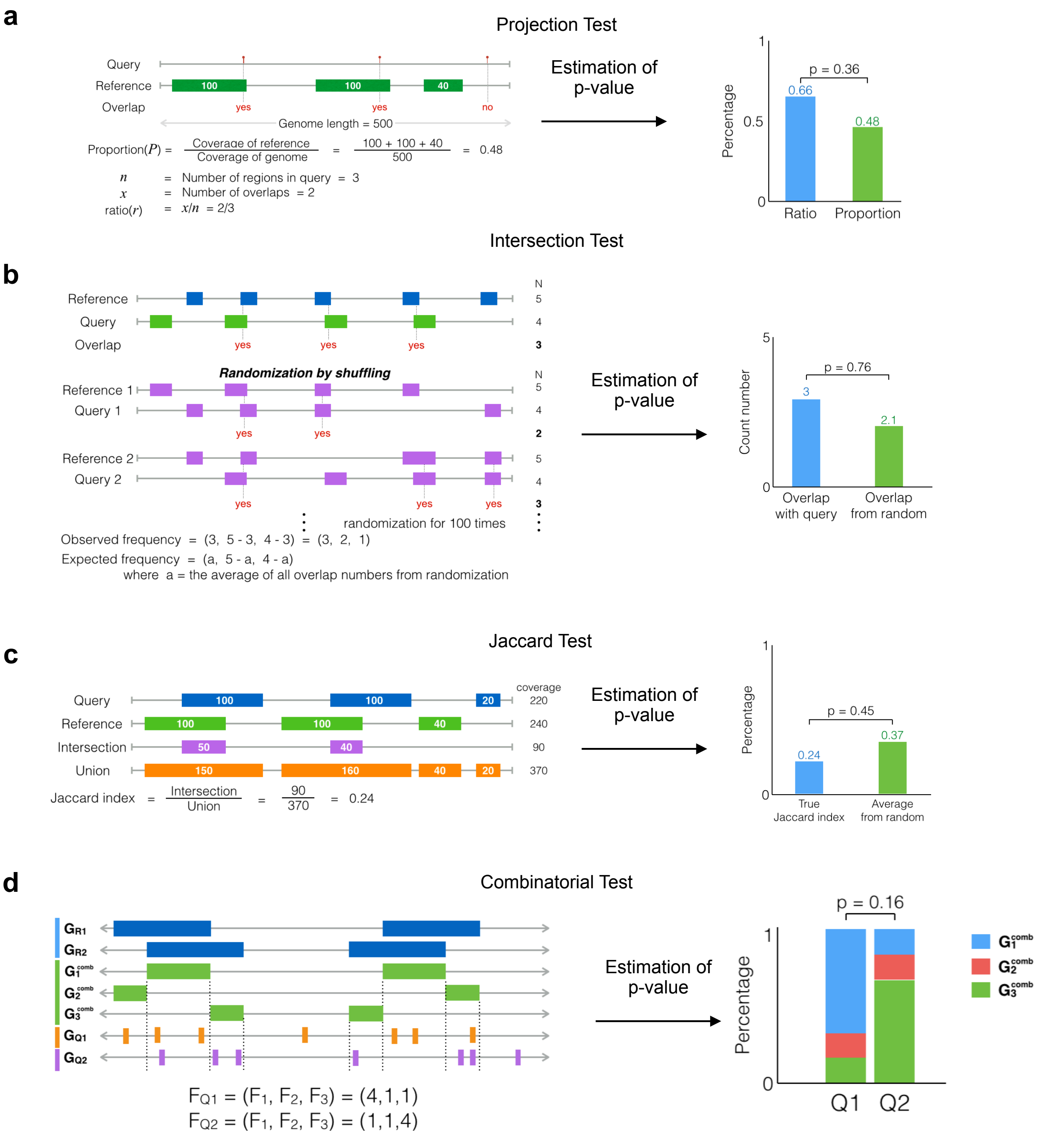


**Supplementary Fig. 2. Schematics of regions versus regions tests in RGT-viz. a,** In the projection test, the query regions are converted into points by their mid-points, then the number of overlaps between reference and point-wise queries are counted. All relevant statistics are calculated such as background proportion (P), which considers the overlap of the reference regions to the original genome, and ratio (r), which considers the overlap between the query and reference region sets. **b,** The intersection test counts the overlaps between query and reference as observed frequency. Randomization is performed by re-sampling region sets from the intersection of both query and reference genomic region sets. This distribution is used to estimate an empirical p-value. **c,** Jaccard test calculates the amount of the overlap, which is the ratio of intersection to the union. Randomization is also performed at least 1000 times. The true Jaccard index is compared with the average of random Jaccard indexes graphically and used to estimate empirical p-values. **d,** The combinatorial test counts the overlaps between each query and each combination of references. These numbers form a profile for each query and can also be compared with one another graphically and using a chi-square test.

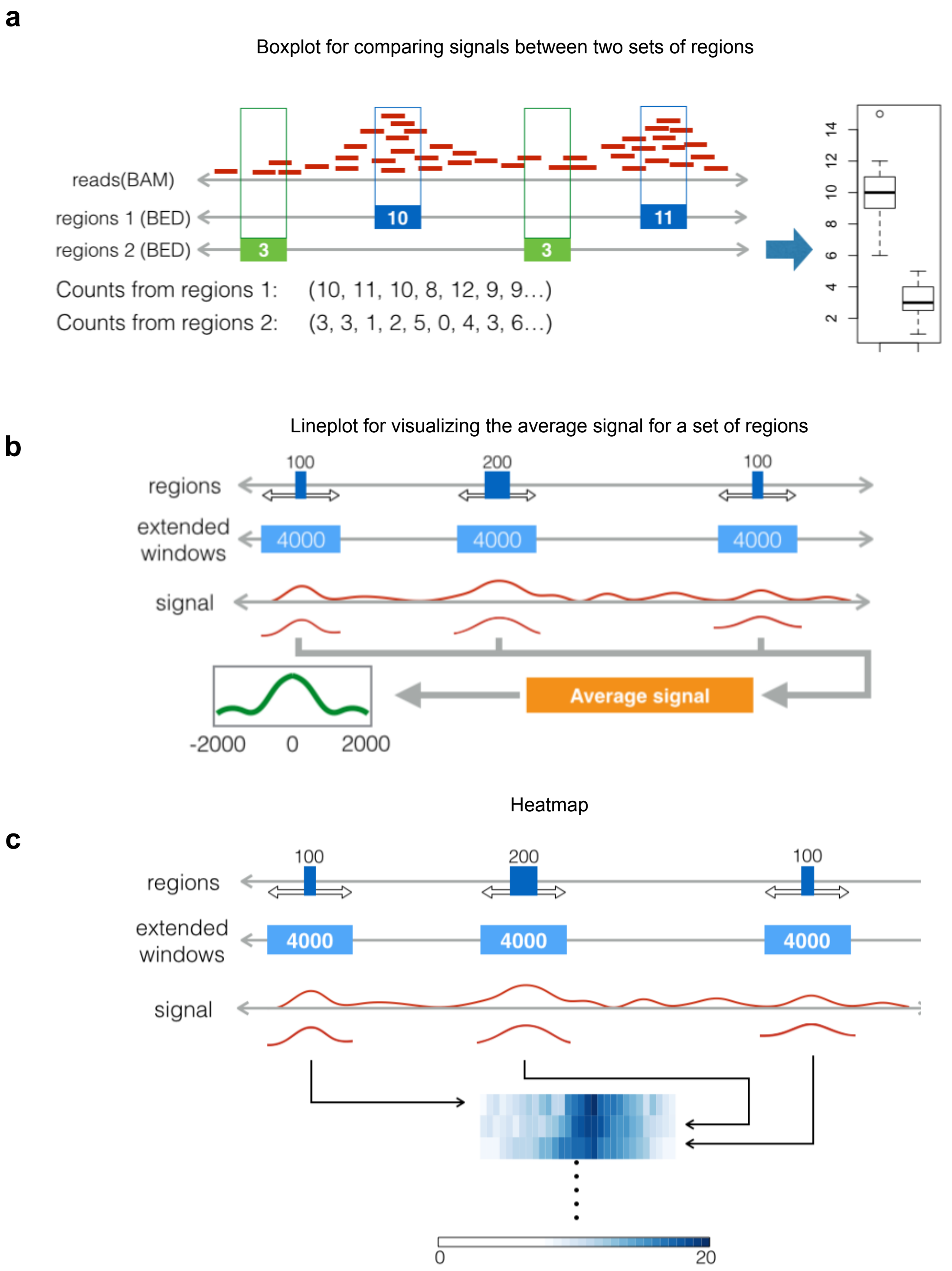


**Supplementary Fig. 3.** Schematics of plotting tools in RGT-viz. **a,** The boxplot compares the reads from one BAM file between two sets of regions represented as BED files. The numbers of reads which fall into the regions of interest are counted, normalized, and compared. **b,** The line plot visualizes the signal of a set of regions. The regions are extended from their midpoint to a window of the same size. According to these windows, the signals on the genome are averaged to form the line plot. **c,** In contrast to the line plot, the heatmap displays all the read counts over the region (x-axis) over distinct regions (y-axis) in a two-dimensional matrix. Colors correspond to the number of reads.

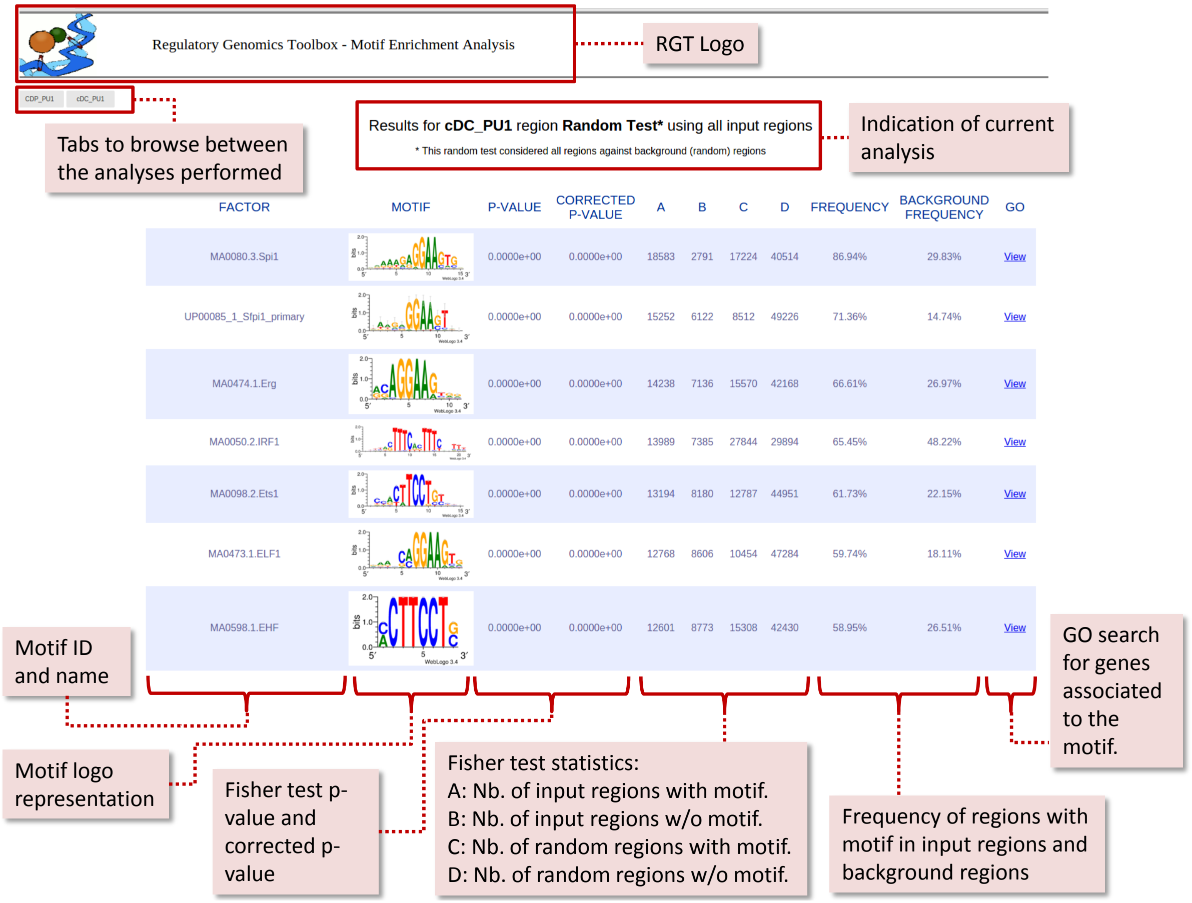

Results for **cDC_PU1** region **Random Test*** using all input regions

* This random test considered all regions against background (random) regions

| FACTOR | MOTIF | P-VALUE | CORRECTED P-VALUE | A | B | C | D | FREQUENCY | BACKGROUND FREQUENCY | GO |
|---|---|---|---|---|---|---|---|---|---|---|
| MA0080.3.Spi1 | | 0.0000e+00 | 0.0000e+00 | 18583 | 2791 | 17224 | 40514 | 86.94% | 29.83% | View |
| UP00085_1_Sfpi1_primary | | 0.0000e+00 | 0.0000e+00 | 15252 | 6122 | 8512 | 49226 | 71.36% | 14.74% | View |
| MA0474.1.Erg | | 0.0000e+00 | 0.0000e+00 | 14238 | 7136 | 15570 | 42168 | 66.61% | 26.97% | View |
| MA0050.2.IRF1 | | 0.0000e+00 | 0.0000e+00 | 13989 | 7385 | 27844 | 29894 | 65.45% | 48.22% | View |
| MA0098.2.Ets1 | | 0.0000e+00 | 0.0000e+00 | 13194 | 8180 | 12787 | 44951 | 61.73% | 22.15% | View |
| MA0473.1.ELF1 | | 0.0000e+00 | 0.0000e+00 | 12768 | 8606 | 10454 | 47284 | 59.74% | 18.11% | View |
| MA0598.1.EHF | | 0.0000e+00 | 0.0000e+00 | 12601 | 8773 | 15308 | 42430 | 58.95% | 26.51% | View |



**Supplementary Fig. 4. An screenshot showing the results of motif enrichment analysis for TF PU.1 (also known as Spi1).** Here, we performed motif matching and enrichment analysis on the PU.1 peaks of the dendritic cell types CDP and cDC. The first two columns showed the motif name and logo found in the repositories. Next, the raw and corrected p-values are shown. The A and B columns represent the number of input regions with at least 1 occurrence of and with no occurrence of that transcription factor. The C and D columns represent the number of background regions with at least 1 occurrence of and with no occurrence of that transcription factor. As we can observe, as expected, PU.1 motifs were ranked in the top (UP00085_1_Sfpi1_primary and MA0080.3.Spi1). Furthermore, we observed that other transcription factors were also enriched in these regions, such as Erg, IRF1, Ets1, ETF1, EHF, and many others. These transcription factors are putative co-binding or regulatory partners of PU.1 and are connected with PU.1 in its regulatory network within the cDC cell type.

Table 1: ChIP-seq data used in the dendritic cell development case study of RGT-viz

| Transcription factor or histone modification | cell type | GEO accession number |
|---|---|---|
| IRF8 | cDC | GSE198651 |
| IRF8 | pDC | GSE198651 |
| PU.1 | MPP | GSE57563 |
| PU.1 | CDP | GSE57563 |
| PU.1 | cDC | GSE57563 |
| PU.1 | pDC | GSE57563 |
| H3K4me1 | MPP | GSE57563 |
| H3K4me1 | CDP | GSE57563 |
| H3K4me1 | cDC | GSE57563 |
| H3K4me1 | pDC | GSE57563 |
| H3K4me3 | MPP | GSE64767 |
| H3K4me3 | CDP | GSE64767 |
| H3K4me3 | cDC | GSE64767 |
| H3K4me3 | pDC | GSE64767 |
| H3K9me3 | MPP | GSE64767 |
| H3K9me3 | CDP | GSE64767 |
| H3K9me3 | cDC | GSE64767 |
| H3K9me3 | pDC | GSE64767 |
| H3K27me3 | MPP | GSE64767 |
| H3K27me3 | CDP | GSE64767 |
| H3K27me3 | cDC | GSE64767 |
| H3K27me3 | pDC | GSE64767 |
| H3K27ac | MPP | GSE73143 |
| H3K27ac | CDP | GSE73143 |
| H3K27ac | cDC | GSE73143 |
| H3K27ac | pDC | GSE73143 |

Table 2: Statistical results of intersection test between PU.1 and IRF8 ChIP-seq peaks across different cell types. Rows are sorted based on $\chi^2$ statistics.

| Ref. name | Query name | Ref. number | Query number | Intersect. | Average intersect. | $\chi^2$-stat. | Positive association p-value | Negative association p-value |
|---|---|---|---|---|---|---|---|---|
| cDC_PU1 | cDC_IRF8 | 20054 | 34003 | 13973 | 6538 | 4489 | 0 | 1 |
| CDP_PU1 | cDC_IRF8 | 20237 | 34003 | 14003 | 6574 | 4438 | 0 | 1 |
| pDC_PU1 | cDC_IRF8 | 21050 | 34003 | 14307 | 6757 | 4367 | 0 | 1 |
| MPP_PU1 | cDC_IRF8 | 6212 | 34003 | 4412 | 1163 | 3359 | 0 | 1 |
| pDC_PU1 | pDC_IRF8 | 21050 | 6467 | 4137 | 1478 | 2100 | 0 | 1 |
| CDP_PU1 | pDC_IRF8 | 20237 | 6467 | 4078 | 1494 | 1979 | 0 | 1 |
| cDC_PU1 | pDC_IRF8 | 20054 | 6467 | 4066 | 1497 | 1955 | 0 | 1 |
| MPP_PU1 | pDC_IRF8 | 6212 | 6467 | 1745 | 848.5 | 351.3 | 5.1e-77 | 1 |
